# Shifts in diversification rates linked to biogeographic movement into new areas: an example of a recent radiation in the Andes

**DOI:** 10.1101/019554

**Authors:** Simon Uribe-Convers, David C. Tank

## Abstract

**Acknowledgements:** We would like to thank S. Mathews for kindly sharing genomic DNA for some taxa used in this study. J. Sullivan, L. Harmon, E. Roalson, J. Beaulieu, B. Moore, N. Nürk, M. Pennell, T. Peterson, and two anonymous reviewers for helpful suggestions or comments on the manuscript. J. Beaulieu, L. Harmon, C. Blair and the University of Idaho Institute for Bioinformatics and Evolutionary Studies (NIH/NCRR P20RR16448 and P20RR016454) for computational aid. Funding for this work was provided by NSF DEB–<u>1210895</u> to DCT for SUC, NSF DEB–<u>1253463</u> to DCT, and Graduate Student Research Grants to SUC from the Botanical Society of America (BSA), the Society of Systematic Biologists (SSB), the American Society of Plant Taxonomists (ASPT), and the University of Idaho Stillinger Herbarium Expedition Funds.

**Premise of the study:** Clade specific bursts in diversification are often associated with the evolution of key innovations. However, in groups with no obvious morphological innovations, observed upticks in diversification rates have also been attributed to the colonization of a new geographic environment. In this study, we explore the systematics, diversification dynamics, and historical biogeography of the plant clade Rhinantheae in the Orobanchaceae, with a special focus on the Andean clade of the genus *Bartsia* L..

**Methods:** We sampled taxa from across Rhinantheae, including a representative sample of Andean *Bartsia* species. Using standard phylogenetic methods, we reconstructed evolutionary relationships, inferred divergence times among the clades of Rhinantheae, elucidated their biogeographic history, and investigated diversification dynamics.

**Key results:** We confirmed that the South American *Bartsia* species form a highly supported monophyletic group. The median crown age of Rhinantheae was determined to be ca. 30 Ma, and Europe played an important role in the biogeographic history of the lineages. South America was first reconstructed in the biogeographic analyses around 9 Ma, and with a median age of 2.59 Ma, this clade shows a significant uptick in diversification.

**Conclusions:** Increased net diversification of the South American clade corresponds with biogeographic movement into the New World. This happened at a time when the Andes were reaching the necessary elevation to host an alpine environment. Although a specific route could not be identified with certainty, we provide plausible hypotheses to how the group colonized the New World.

---

The investigation of global patterns of biodiversity has a long history (e.g., Mittelbach et al., 2007). With the increase in our knowledge of phylogenetic relationships as well as methods for using phylogenies to understand diversification rates and biogeographic patterns (e.g., Ree et al., 2005; Alfaro et al., 2009), these global patterns can now be placed in an explicitly historical context (sensu Moore and Donoghue, 2007). Along these lines, differences in species richness between geographic areas have often been explained by climatic stability, age of the region, and/or niche conservatism that contributes to the slow, but steady, accumulation of species over time (Wiens and Donoghue, 2004). Likewise, clade specific bursts in net diversification (speciation minus extinction) are often associated with the evolution of novel morphologies, referred to as key innovations, such as nectar spurs in angiosperms (e.g., Hodges, 1997), molar characters in mammals (Woodburne et al., 2003), and feathers in birds (Ostrom, 1979). More recently, Moore and Donoghue (2007) demonstrated that in the plant families Adoxaceae and Valerianaceae, shifts in diversification rates were not correlated with the evolution of novel floral characters, but rather, with the movement into new geographic areas, and hypothesized that “dispersification” (dispersal and diversification) may play a larger role in shaping global biodiversity patterns than previously recognized. This is equivalent to the “key opportunity” of Donoghue and Sanderson (2015), and follows the hypothesis that in newly emerging environments, as long as the corridors for biogeographic movements are in place, these new areas will often be filled with lineages from environmentally similar areas where the relevant morphological and/or physiological adaptations are already in place (Donoghue, 2008). Empirical tests of these hypotheses not only require a robust estimate of phylogenetic relationships, but also the estimation of divergence times, diversification rates, and biogeographic patterns for the group of interest.

Various approaches have been taken to assess phylogenetic relationships, divergence times, and rates of diversification – each increasing our understanding of biodiversity and the way in which it has been produced. Bayesian analyses are now regularly used to estimate divergence times (e.g., Bacon et al., 2012; Drummond et al., 2012), most often performed in the program BEAST (Drummond and Rambaut, 2007), because the use of probabilistic priors accommodates both phylogenetic uncertainty (i.e., topology and branch lengths), as well as the timing of calibration points. Diversification rate analyses have been instrumental to our understanding of disparities in clade richnesses across the tree of life. For example, Alfaro et al. (2009) suggested that six pulses of accelerated diversification and three slowdowns, instead of single events, have shaped the current diversity of jawed vertebrates. Likewise, Tank et al. (2015) revealed a pattern of “nested radiations” across angiosperms–where clade-specific upticks in net diversification were nested within earlier diversification rate increases—and suggested that widely heterogeneous diversification rates have generated this pattern. Additionally, in the plant genus *Asclepias* L. (milkweeds) it has been shown that increases in the rate of diversification are tightly associated with the evolution of defense traits that prevent or minimize herbivory, and that this resulted in an adaptive radiation in the group (Agrawal et al., 2009). Moreover, studies of Andean plants, e.g., the family Valerianaceae (Bell and Donoghue, 2005a) and the genus *Lupinus* L. (Hughes and Eastwood, 2006; Drummond et al., 2012; Hughes and Atchison, 2015), have shown that groups with North American temperate ancestors have elevated diversification rates in the Andes, given that they were “pre–adapted” to the conditions of the newly and unoccupied niche at the time that they colonized the Andes (see also the ‘key landscape’ concept of Givnish [1997; 2015] to explain increased diversification rates in montane regions).

To investigate the influence of biogeographic movements on rates of diversification, we have chosen to study the mostly European clade Rhinantheae of the parasitic plant family Orobanchaceae (Wolfe et al., 2005; Bennett and Mathews, 2006; McNeal et al., 2013), with a particular focus on the genus *Bellardia* All., a clade of 48 species that are distributed across two disjunct geographic regions. The majority of the species in *Bellardia* were formerly part of the genus *Bartsia* L., which has been recently recircumscribed (Scheunert et al., 2012) to better reflect the evolutionary history of its species. Prior to this taxonomic rearrangement, *Bartsia* (49 spp.) had two species distributed in the mountains of northeastern Africa (*B. decurva* Benth. and *B. longiflora* Benth.), one in the Mediterranean region (*B. trixago* L.), one in Scandinavia, the Alps, Greenland and the Hudson Bay region of northeastern North America (*B. alpina* L.), and the remaining 45 species distributed throughout the páramos of Andean South America (Molau, 1990). Broad–scale phylogenetic studies of Orobanchaceae (Wolfe et al., 2005; Bennett and Mathews, 2006) and the Rhinantheae clade (Těšitel et al., 2010) had suggested that *Bartsia* was not monophyletic, but Scheunert et al. (2012) were the first to include species from the complete geographic distribution of the genus, as well as the two species of the related Mediterranean genus *Parentucellia* Viv.. However, because their sampling only included two species of the South American clade of *Bartsia*, they chose to only reclassify these two species, leaving ca. 43 species in a large polyphyletic group with the monotypic lineage of *B. alpina* in Europe. The South American *Bartsia* species, which we will refer to here as the Neobartsia clade, are quite distinct from their Mediterranean counterparts (i.e., *Bellardia trixago,* and the two species of *Parentucellia* that were also moved to the expanded genus—*Bellardia latifolia* and *B. viscosa*) in multiple aspects. Ecologically, species in the Neobartsia clade grow at high elevation (ca. 3,000–5000 m) in wet environments while the Mediterranean species grow at low elevation (ca. 0–500 m) in seasonally dry environments. Geographically, the Neobartsia clade is restricted to the Andes while the Mediterranean taxa are native to the Mediterranean region and more recently have been introduced to Australia, coastal Chile, and coastal western North America. Finally, the Mediterranean species all have reflexed corolla lips, usually associated with bee pollination, whereas a large number of the species in the Neobartsia clade have erect corolla lips that are thought to be associated with hummingbird pollination due to their tubular shape and the placement of reproductive parts.

Previous studies of the group have only included a minor fraction of the South American species richness, usually sampling only one or two species, making it difficult to assess the influence of biogeographic movements on rates of diversification across the clade. Here, we included representatives from all clades (sensu Molau, 1990) comprising the former genus *Bartsia,* including a morphologically and geographically representative sampling of the South American diversity, as well as a representative sampling of all known allied genera of the Rhinantheae clade of Orobanchaceae, to establish a robust and well–supported phylogeny of the clade based on both chloroplast and nuclear ribosomal DNA sequence data. We then use this phylogeny to estimate divergence times across the clade and to investigate the biogeographic history of the clade, with a special focus on the origin of the Neobartsia clade in Andean South America. Finally, we use all these analyses to test if increases in rates of diversification are indeed associated with biogeographic movements into newly formed environments, i.e., “dispersification” sensu Moore and Donoghue (2007).

## MATERIALS AND METHODS

### Sampling

A total of 49 taxa were included in this study (Table 1), with newly collected specimens stored in airtight plastic bags filled with silica gel desiccant in the field. Because our main focus is the diversification dynamics of the South American Neobartsia clade in the context of the disparate geographic distributions of the Old World species and the remainder of the mostly European Rhinantheae clade of Orobanchaceae, our sampling effort included representatives of 10 of the 11 genera thought to comprise the clade (Wolfe et al., 2005; Bennett and Mathews, 2006; Těšitel et al., 2010; Scheunert et al., 2012; McNeal et al., 2013). *Bellardia* and the Neobartsia clade are represented here by 15 South American species and two of the three Mediterranean taxa, *Bellardia trixago* (L.) All. and *Bellardia viscosa* (L.) Fisch. & C.A. Mey. The South American species were selected to encompass the morphological and geographic diversity in the clade. Based on previous results (Olmstead et al., 2001; Wolfe et al., 2005; Bennett and Mathews, 2006; Těšitel et al., 2010; Scheunert et al., 2012; McNeal et al., 2013) *Melampyrum* L. was used as the outgroup for the Rhinantheae clade.

**Table 1.** Taxa and voucher information for plant material from which DNA was extracted. s.n = *sine numero* (without a collecting number). Herbarium abbreviations are as follow: FHO = University of Oxford Herbarium, K = Royal Botanic Gardens Kew, ID, University of Idaho = Stillinger Herbarium, ANDES = Museo de Historia Natural Universidad de los Andes, WTU = University of Washington Herbarium, GH = Harvard University Herbarium, LJU = University of Ljubljana Herbarium, USFS = United States Forest Service, CBFS = University of South Bohemia České Budějovice. GenBank accessions for sequences not generated in this study are also shown.

| Species | DNA Voucher/Herbarium | GenBank Accession Number |  |  |  |  |
| --- | --- | --- | --- | --- | --- | --- |
|  |  | ITS | ETS | trnT-trnL | trnL-trnF | rps16 |
| *Bartsia alpina* L. | Lampinen s.n/ID | FJ790046 | KM408206 | KM408239 | KM434119 | N/A |
| *B. crenoloba* Wedd. | Solomon 7152/K | KM408228 | KM408185 | KM408240 | KM434106 | KM408308 |
| *B. laniflora* Benth. | SU-24/ANDES | KM408221 | KM408174 | KM408242 | KM434110 | KM408307 |
| *B. laticrenata* Benth. | Ramsay & Merrow-Smith 771/K | KM408219 | KM408178 | KM408243 | KM434104 | N/A |
| *B. melampyroides* (Kunth) Benth. | Tank 2005-07/WTU | KM408218 | KM408186 | KM408245 | KM434114 | KM408300 |
| *B. orthocarpiflora* Benth. | Ollgaard 34129/K | KM408216 | KM408184 | KM408246 | KM434111 | KM408296 |
| *B. pedicularoides* Benth. | Jorgenson 1729/K | FJ790047 | N/A | FJ790077 | N/A | N/A |
| *B. pyricarpa* Molau | Tank 2005-36/WTU | KM408226 | KM408182 | KM408247 | KM434102 | KM408294 |
| *B. ramosa* Molau | CG-016/ANDES | KM408229 | KM408176 | KM408248 | KM434108 | KM408304 |
| *B. santolinifolia* (Kunth) Benth. | SU-18/ANDES | KM408220 | KM408175 | KM408249 | KM434109 | KM408306 |
| *B. sericea* Molau | Tank 2005-06/WTU | KM408224 | KM408181 | KM408250 | KM434105 | KM408297 |
| *B. cf sericea* Molau | Tank 2005-25/WTU | KM408227 | KM408179 | KM408251 | KM434101 | KM408302 |
| *B. cf inaequalis* Benth. *ssp. duripilis* (Edwin) Molau | Tank 2005-29/WTU | KM408225 | KM408183 | KM408252 | KM434107 | KM408295 |
| *B. stricta* (Kunth) Benth. | SU-1b/ANDES | KM408222 | KM408177 | KM408253 | KM434112 | KM408305 |
| *B. tenuis* Molau | Tank 2005-02/WTU | KM408223 | KM408180 | KM408254 | KM434103 | KM408299 |
| *B. thiantha* Diels | RGO 2009-23/WTU | KM408217 | KM408187 | KM408255 | KM434113 | KM408303 |
| *Bellardia trixago* (L.) All. | Bennett s.n/FHO | FJ790063 | KM408189 | KM408256 | KM434100 | KM408301 |
| *B. viscosa* (L.) Fisch. & C.A. Mey | Halse 2249/ID | AY911244 | KM408188 | KM408273 | KM434095 | KM408298 |
| *Bornmuellerantha aucheri* (Boiss.) Rothm. | Oganesian et al. 03-1575/K | KM408237 | KM408197 | KM408267 | KM434116 | KM408279 |
| *Euphrasia alsa* F.Muell. | Zich 220/GH | KM408212 | KM408202 | KM408257 | KM434084 | KM408282 |
| *E. collina* R.Br. | Zich 209/GH | N/A | KM408203 | KM408258 | KM434086 | KM408284 |
| *E. mollis* (Ledeb.) Wettst. | Mancuso 107/ID | KM408213 | N/A | KM408259 | KM434082 | KM408281 |
| *E. regelii* Wettst. | Ho 1741/GH | KM408214 | N/A | KM408260 | KM434085 | KM408285 |
| *E. stricta* D. Wolff ex J.F. Lehm | Musselman 4872/ID | KM408215 | N/A | KM408261 | KM434098 | KM408283 |
| *Hedbergia abyssinica* (Benth.) Molau var. *abyssinica* | Etuge 3488/K | FJ790061 | KM408194 | KM408262 | KM434120 | KM408291 |
| *H. abyssinica* (Benth.) Molau var. *nykiensis* | Carter et al 2386/K | KM408231 | KM408193 | KM408263 | KM434122 | N/A |
| *H. abyssinica* (Benth.) Molau var. *petitiana* | Paton s.n/K | KM408230 | KM408195 | KM408264 | KM434123 | KM408290 |
| *H. decurva* (Hochst. ex Benth.) A. Fleischm. & Heubl | Wesche 9/K | N/A | KM408191 | KM408241 | KM434121 | KM408292 |
| *H. longiflora* ssp. *longiflora* (Hochst. ex Benth.) A. Fleischm. & Heubl | Kisalye van Heist 109/K | KM408232 | KM408192 | KM408244 | KM434099 | KM408286 |
| *Lathraea squamaria* L. | Frajman s.n/LJU | FJ790044 | KM408204 | EU264174 | KM434087 | KM408309 |
| *Melampyrum carstiense* Fritsch | Krajsek s.n/LJU | GU445314 | N/A | EU264177 | KM434088 | KM408315 |
| *M. lineare* Lam. | Bjork 6465/ID | KM408208 | KM408207 | KM408265 | KM434096 | KM408316 |
| *M. sylvaticum* L. | Krajsek s.n/LJU | EU624134 | N/A | KM408266 | KM434089 | KM408314 |
| *Odontites corsicus* (Loisel.) G. Don | J. Stefani/ID | KM408238 | KM408200 | KM408268 | KM434117 | N/A |
| *O. linkii* Heldr. & Sart. ex Boiss. ssp. *cyprius* | Ferguson 4537/K | KM408234 | KM408196 | KM408269 | KM434083 | KM408288 |
| *O. maroccanus* Bolliger | Gattefose s.n/K | KM408233 | KM408198 | KM408270 | KM434097 | N/A |
| *O. vulcanicus* Bolliger | Bolliger & Moser O-M3/K | KM408235 | KM408199 | KM408271 | KM434115 | KM408289 |
| *O. vulgaris* Moench | Kharkevich s.n/K | KM408236 | KM408201 | KM408272 | KM434118 | KM408287 |
| *Rhinanthus crista-galli* L. | Bjork 6656/ID | KM408210 | N/A | KM408274 | KM434091 | KM408313 |
| *R. freynii* (A.Kern. ex Sterneck) Fiori | Mathews 04-05 | GU445319 | KM408205 | KM408275 | KM434092 | KM408310 |
| *R. kyrollae* Chabert | Stickney 1236/USFS | KM408209 | N/A | KM408276 | KM434090 | KM408312 |
| *R. serotinus* (Schönh.) Oborny | Musselman 4871/ID | KM408211 | N/A | KM408277 | KM434093 | KM408311 |
| *Rhynchocorys elephas* Griseb. | Tesitel 5044/CBFS | FJ790055 | N/A | FJ790085 | N/A | N/A |
| *R. kurdica* Nábělek | Tesitel 5042/CBFS | FJ790037 | N/A | FJ790067 | N/A | N/A |
| *R. maxima* Richter | Tesitel 5040/CBFS | FJ790037 | N/A | FJ790067 | N/A | N/A |
| *R. odontophylla* R.B.Burbidge & I.Richardson | Tesitel 5038/CBFS | FJ790034 | N/A | FJ790064 | N/A | N/A |
| *R. orientalis* Benth. | Tesitel 5039/CBFS | FJ790035 | N/A | FJ790065 | N/A | N/A |
| *R. stricta* Albov | Tesitel 5047/ CBFS | FJ790057 | N/A | FJ790087 | N/A | N/A |
| *Tozzia alpina* L. | Mathews 04-04 | AY911258 | KM408190 | KM408278 | KM434094 | KM408280 |

### Molecular Methods

Total genomic DNA was extracted from silica gel–dried tissue or herbarium material using a modified 2X CTAB method (Doyle and Doyle, 1987). Two chloroplast (cp) regions–*trnT*–*trnF* region and the *rps16* intron–were amplified via polymerase chain reaction (PCR) using the *trn*–a and *trn*–f (Taberlet et al., 1991) and the *rps16_*F and *rps16_*R primers (Oxelman et al., 1997), respectively. The nuclear ribosomal (nr) internal transcribed spacer (ITS) and external transcribed spacer (ETS) regions were amplified using the ITS4 and ITS5 primers (Baldwin, 1992) and the ETS–B (Beardsley and Olmstead, 2002) and 18S–IGS (Baldwin and Markos, 1998), respectively. PCR profiles for all regions followed Tank and Olmstead (2008). When amplification of a region in one fragment was not possible, internal primers were used to amplify the region in multiple fragments. The primer pairs *trn–a/trnb, trnc/trn–d,* and *trn–e/trn–f* (Taberlet et al., 1991) were used to amplify the *trnT*–*trnF* region. Additionally, *Bellardia* specific internal primers were designed and used when these primer combinations failed (*trnT/trnL* intergenic spacer: *trnT–L_iF* 5–CTTGGTTTTCATCCGTAAAGG–3 and *trnT–L_iR* 5–CCTTTACGGATGAAAACCAAG–3). Following Tank and Olmstead (2008), the *rps16*_F/*rps16*_iR and *rps16*_iF/*rps16*_2R primer combinations were used to amplify the *rps16* intron in two fragments. Similarly, the ITS5/ITS2 and ITS3/ITS4 primer combinations (Baldwin, 1992) were used to amplify the ITS region in two fragments.

PCR products were purified by precipitation in a 20% polyethylene glycol 8000 (PEG)/2.5 M NaCl solution and washed in 70% ethanol prior to sequencing. To ensure accuracy, we sequenced both strands of the cleaned PCR products on an ABI 3130xl capillary DNA sequencer (Applied Biosystems, Foster City, California, USA) using ABI BigDye v.3.1 cycle sequencing chemistry. Sequence data were edited and assembled for each region using the program Sequencher v.4.7 (Gene Codes Corp., Ann Arbor, Michigan, USA), and consensus sequences were generated and submitted to GenBank (GenBank accessions: ETS: KM408174–KM408207, ITS: KM408208–KM408238, *trnT–trnL*: KM408239–KM408278, *rps16*: KM408279–KM408316, *trnL–trnF*: KM434082–KM434123). When sequencing was not possible for any given species or gene region, GenBank sequences were used to reduce the amount of missing data in the final matrix (Table 1).

### Phylogenetic Analyses

Although two separate cpDNA regions were sequenced, the evolutionary histories of the *trnT–trnF* region and the *rps16* intron are tightly linked due to the nonrecombining nature of the chloroplast genome, and thus, were treated as a single locus. The nrDNA regions (ITS and ETS) were also treated as a single locus, given that they are linked because of their physical proximity in the nrDNA repeat. We created three primary datasets with our two independent loci: 1) cpDNA only, 2) nrDNA only, and 3) a combined cpDNA and nrDNA dataset. Given the wide phylogenetic diversity of sampled taxa, and relatively high molecular evolutionary rates of the nrDNA and cpDNA regions employed here, global alignments using default settings in Muscle v.3.6 contained many ambiguously aligned regions. Therefore, global alignments across the Rhinantheae clade were created for each gene region using the group–to–group profile alignment method as implemented in Muscle v.3.6 (Edgar, 2004). The group–to–group profile alignment method takes advantage of previous knowledge about monophyly of the major lineages (e.g., Těšitel et al., 2010; Scheunert et al., 2012) and consists of lineage–specific alignments that are then iteratively aligned to one another resulting in fewer alignment ambiguities (Smith et al., 2009). These alignments were visually inspected and minor adjustments were made manually using Se–Al v.2.0a11 (Rambaut, 1996). Sites that could not be unambiguously aligned were excluded from the analyses. File format conversions and matrix concatenations were performed using the program Phyutility v.2.2 (Smith and Dunn, 2008).

A statistical selection of the best–fit model of nucleotide substitution according to the Akaike information criterion (AIC) was conducted independently for each gene region using the program jModelTest (Guindon and Gascuel, 2003; Posada, 2008). Based on these results, partitioned (by gene region) maximum likelihood (ML) analyses were performed on our three primary datasets using RAxML v. 7.2.4 (Stamatakis, 2006) with 1,000 replicates of nonparametric bootstrapping using the rapid bootstrap algorithm (Stamatakis et al., 2008). Every fifth bootstrap tree generated by the rapid bootstrap analyses was used as a starting tree for full ML searches and the trees with the highest ML scores were chosen. Likewise, partitioned Bayesian inference (BI) analyses were performed using the parallel version of MrBayes v. 3.1.2 (Ronquist and Huelsenbeck, 2003) with the individual parameters unlinked across the data partitions. Analyses consisted of two independent runs with four Markov chains using default priors and heating values. Each independent run consisted of 15 million generations and was started from a randomly generated tree and was sampled every 1,000 generations. Convergence of the chains was determined by analyzing the plots of all parameters and the –lnL using Tracer v.1.5 (Rambaut and Drummond, 2004). Stationarity was assumed when all parameters values and the –lnL had stabilized; the likelihoods of independent runs were considered indistinguishable when the average standard deviation of split frequencies was < 0.001. Consensus trees were obtained for each dataset using the sumt command in MrBayes. Finally, incongruences between the cpDNA and the nrDNA topologies were investigated using the approximately unbiased (AU) test (Shimodaira, 2002) and the Shimodaira–Hasegawa (SH) test (Shimodaira and Hasegawa, 1999), as implemented in the program CONSEL (Shimodaira and Hasegawa, 2001).

### Divergence Time Estimation

To maximize the number of taxa in our dating analyses and to improve branch length estimation by minimizing the amount of missing data (Lemmon et al., 2009), we reduced our combined dataset to include sequences for only the cpDNA *trnT–trnL* intergenic spacer and the nrDNA ITS region. This resulted in a dataset that included all 49 taxa and only 2% missing data, compared to 15% missing data for the complete dataset. Each gene was treated as a separate partition. To ensure convergence in divergence times, five independent runs were conducted using BEAST v.1.5.4 (Drummond and Rambaut, 2007). BEAST implements Markov Chain Monte Carlo (MCMC) methods that allow for uncertainty in both the topology and the calibration points, i.e., calibration points are treated as probabilistic priors, rather than point estimates (Ho and Phillips, 2009). It also implements an uncorrelated lognormal relaxed clock (UCLN) (Drummond et al., 2006), allowing every branch to have an independent substitution rate.

Each run was started from the resulting ML tree obtained for the dataset containing all regions, after performing a semiparametric rate smoothing based on penalized likelihood (Sanderson, 2002) in R (R Development Core Team, 2013) using the package Ape (Paradis et al., 2004). Each run consisted of 100,000,000 generations sampled every 1000 trees. The models of nucleotide substitution were kept unlinked for both partitions and the tree priors were kept as default under the birth–death process.

Because of the mostly herbaceous habit of the species in Orobanchaceae, there are no known fossils for the family. This lack of fossils made the dating of our analyses dependent on secondary calibrations obtained from a previous study. Based on explicit age estimates of an ITS molecular clock (Wolfe et al., 2005), a calibration point at the node containing every genus except *Melampyrum* (i.e., one node younger than the root) was used. This was done with a lognormal distribution prior with an offset of 25 million years (Ma), a mean of 0.9, and a standard deviation of 0.8, this way incorporating uncertainty in the calibration point. Because the use of this secondary calibration is far from ideal, to corroborate our calibration strategy, an additional analysis using the most recent uplift of the Andes as the calibration point (Simpson, 1975; Burnham and Graham, 1999; Gregory-Wodzicki, 2000; Antonelli et al., 2009) was conducted with a lognormal distribution prior (offset of 1.7 Myr, a mean of 0.2 and an standard deviation of 0.6). This calibration prior was set at the node where the species in the Neobartsia clade diverge from *B. viscosa*. This additional calibration scenario was conducted to assess the impact of alternative calibration points in the node ages.

Convergence of the parameters was monitored using Tracer v. 1.5 and the resulting trees were summarized using TreeAnnotator v.1.5.4 (Drummond and Rambaut, 2007) after 25% of the trees had been discarded as burn–in. Each of the five topologies and their node heights were visualized using FigTree v. 1.3.1 (Rambaut, 2006) and a final tree, representing the maximum clade credibility tree with information of the 95 percent highest posterior density (HPD), was obtained by combining the five runs using LogCombiner v.1.5.4 (Drummond and Rambaut, 2007) and by summarizing them with TreeAnnotator v.1.5.4.

### Biogeographic Analyses

The biogeographic history of *Bellardia* and allied genera was reconstructed using the program Lagrange v. C++ (Ree and Smith, 2008). Lagrange implements the maximum likelihood Dispersal–Extinction–Cladogenesis (DEC) model (Ree et al., 2005) to estimate the most likely ancestral geographic range based on current distributions of extant lineages. The DEC model has been used broadly to study biogeographic patterns in closely related taxa with restricted geographic areas (e.g., Fabre et al., 2013), large genera with cosmopolitan distributions (e.g., Emadzade et al., 2011), intercontinental migrations within families (e.g., Clayton et al., 2009), and large, diverse clades of angiosperms (e.g., Beaulieu et al., 2013). This model assumes extinction or dispersal by contraction or expansion of the ancestral geographic range, respectively. Not only does Lagrange find the most likely ancestral area at a node, it calculates the probability of that area and compares it to other competing biogeographic scenarios. One of the caveats of this method, however, is that parameter space increases rapidly with the addition of geographic areas—it is currently advised to maintain the number of geographic areas for an unconstrained model to seven or eight. To alleviate this potential problem and introduce an additional advantage to biogeographic analyses, the user is given the option to assign a dispersal probability matrix based on prior knowledge of connectivity between areas, incorporating valuable ancestral geographic information. This is particularly important when proposed geographic areas are available only at certain time periods for various reasons—e.g., the formation or absence of a land bridge between continents, the uplift of a mountain, the formation of an island, etc.—which can result in more realistic dispersal routes. In plants however, most of this knowledge, at least for Northern Hemisphere temperate plants, is based on macrofossils of woody mesophytic taxa, e.g., *Quercus* (Tiffney and Manchester, 2001). Because the herbaceous genus *Bellardia* is almost completely restricted to alpine–like conditions that separate this lineage from the ecological conditions in which mesophytic forest species are found, and the vast majority of the Rhinantheae clade is also herbaceous, we consider that biological routes for these types of taxa are less well understood (Donoghue and Smith, 2004), and therefore, we did not include a dispersal probability matrix in our analyses (i.e., equal transition rates between all areas; see also Smith and Donoghue, 2010).

We used Lagrange on a posterior distribution of 1,000 randomly chosen trees (post burn–in) from our dating analyses. By inferring ancestral ranges over a posterior distribution of trees we are incorporating uncertainty in both topology as well as times of divergence (Smith, 2009; Smith and Donoghue, 2010; Beaulieu et al., 2013). We conducted three independent analyses with varying distributions of current taxa. The first analysis was performed with conservative geographic ranges following Mabberley’s Plant–Book (Mabberley, 2008), in which the genera have wider distributions, e.g,. the genus *Euphrasia* L. has a north temperate distribution (Eurasia, Europe and Eastern North America). The second analysis included prior expert knowledge about the distribution of the genera based on published work, e.g., we followed the proposed Eurasian origin for the genus *Euphrasia* (Gussarova et al., 2008). The final analysis was based on species–specific distributions based on the explicit species that we sampled, i.e., species within a genus can have different distributions to account for endemisms and/or disparate distributions within a genus.

We considered species to be distributed in five distinct geographic areas: i) Eurasia (western Eurasia: the Balkan Peninsula and the Caucasus region), ii) Europe (including the Mediterranean climatic region in southern Europe and northern Africa), iii) Africa (montane northeastern Africa), iv) North America (Hudson Bay region of northeastern North America), and v) South America (including only the Andes). The results of the analyses were summarized in R. Following Beaulieu et al. (2013), we calculated Akaike weights for every biogeographic scenario reconstructed at every node in each tree separately. We then summed the Akaike weights for each node and averaged them across the distribution of trees, which resulted in composite Akaike weights (*w*_*i*_) for our biogeographic reconstructions. This means that the composite Akaike weights (*w*_*i*_) can be interpreted as being the relative likelihood of a given biogeographic scenario compared to other possible scenarios. Furthermore, we examined the evidence for the most supported scenario by calculating an evidence ratio of this model versus the next most supported model (Burnham and Anderson, 2002), i.e. the evidence ratio is the relative likelihood of one model versus another (*w*_*i*_/*w*_*j*_). These were interpreted as relative evidence of one scenario being the most supported when comparing it against competing biogeographic hypotheses (Beaulieu et al., 2013).

### Diversification Rates

Diversification rate analyses were conducted on the same posterior distribution of 1,000 trees, as well as on the maximum clade credibility (MCC) tree using MEDUSA (Alfaro et al., 2009), which is an extension of the method described by Rabosky et al. (2007) and is available in the R package geiger 2.0 (Pennell et al., 2014). In Rabosky et al. (2007), two likelihoods are estimated for a dated tree: i) a phylogenetic likelihood that uses the timing of the splits on the backbone to estimate ML values for birth and death rates following the equations of Nee et al. (1994), and ii) a taxonomic likelihood that uses species richness along with the date of the splits, estimating diversification rates following Magallón and Sanderson (2001). MEDUSA (Alfaro et al., 2009) looks for shifts in diversification rates in a stepwise manner by comparing AIC scores of successively more complex models. This method requires complete sampling that is achieved by collapsing clades of interest to a single tip and then assigning clade richnesses to these tip lineages. We collapsed our trees into tips representing each of the major lineages of Rhinantheae, which in most cases corresponded to each of the genera. The following clade richnesses were used: Neobartsia clade (45 spp.), *Parentucellia* clade (2 spp.), *Bellardia* clade (1 sp.), *Odontites* clade (32 spp.), *Euphrasia* clade (350 spp.), *Rhinanthus* clade (45 spp.), *Melampyrum* clade (35 spp.), *Lathraea* clade (7 spp.), *Rhynchocorys orientalis* (1 spp.), *Rhynchocorys elephas* (1 spp.), *Rhynchocorys odontophylla* (1 spp.), *Rhynchocorys kurdica* (1 spp.), *Rhynchocorys stricta* (1 spp.), *Rhynchocorys maxima* (1 spp.), *Bartsia alpina* (1 spp.), *Tozzia alpina* (1 spp.), *Hedbergia abyssinica* (1 spp.), *Hedbergia decurva* (1 spp.), *Hedbergia longiflora* (1 spp.).

To compare our MEDUSA results we conducted an additional analysis using the program SymmeTREE v1.0 (Chan and Moore, 2005). SymmeTREE is based on the topological distribution of species on the whole tree, which is compared to a distribution simulated on a tree under the equal–rates Markov random branching model (EMR), where the probability of a branching event is constant throughout the tree (Yule, 1924). If a clade shows an unbalanced distribution of species richness when compared to its sister clade, then a shift in the rate of diversification is identified. SymmeTREE also estimates several whole tree statistics that are evaluated against their own simulated null distribution, i.e., a constant pure–birth model (Chan and Moore, 2005). To accommodate topological and temporal uncertainty, we assessed diversification rate shifts with SymmeTREE using default settings across a random set of 542 trees from the posterior distribution of trees from our divergence time analysis; the full set of 1,000 trees was not used due to computational limitations.

## RESULTS

### Molecular Methods

The cpDNA data set included the *trnT–trnF* region and the *rps16* intron and had a total length of 2,686 bp with 13% missing data. Similarly, the nrDNA dataset included the ITS and ETS regions with a total of 1,134 bp and 17% missing data. A combined data set was created from the cpDNA and the nrDNA matrices, with a total length of 3,820 bp and 15% missing data (files deposited in the Dryad Digital Repository: *data will be submitted after acceptance of the manuscript*).

### Phylogenetic Analyses

Alignment of individual gene regions was straightforward requiring minor adjustments to the automated alignment strategy implemented in MUSCLE v. 3.6 (matrices and trees are available on TreeBase: *Temporary reviewer access* http://purl.org/phylo/treebase/phylows/study/TB2:S11528?x–access–code=334b76effd95e3f56eb4ffe0185fc9ad&format=html). Some regions that could not be unambiguously aligned in the *trnT–trnL* intergenic spacer and in the ETS region were excluded from the analyses (*trnT–L*: alignment positions 519–529 and 587–620; ETS: alignment positions 63–65, 83–85 and 152–158). Model selection for the cpDNA regions yielded the General Time Reversible model + Γ (GTR) (Rodríguez et al., 1990) for the *trnT–trnF* intergenic spacer, and the Transversion model + Γ (TVM) for the *rps16* intron. The ITS and ETS regions resulted in the selection of the GTR+I+Γ and Hasegawa–Kishino–Yano+ Γ (HKY) models, respectively. To avoid the difficulties of estimating Γ and the invariable sites simultaneously (Ronquist and Huelsenbeck, 2003; Yang, 2006), the model of substitution GTR+Γ with an increase in the number of rate categories from four to six was preferred in the case of the ITS region.

Our results from every dataset (Fig. 1 for the combined dataset and Fig. 2 for the cpDNA and nrDNA datasets) are in concordance with those presented in previous Rhinantheae studies (Těšitel et al., 2010; Scheunert et al., 2012), and assessment of incongruences between the cpDNA and nrDNA datasets showed that these were either not significant, or if they were, the alternative topology was only weakly supported. For example, the well–supported relationships in the cpDNA dataset between *Tozzia alpina* and *Hedbergia*—or between *Odontites* and *Bellardia*—are not statistically significant when constrained in the nrDNA dataset. Conversely, the relationship between *H. abyssinica* var. *nykiensis* and *H. decurva* found in the cpDNA dataset is significant in the AU Test, but it is only moderately supported on the tree (BS 72, PP 0.96) and it does not exist in the combined dataset (Table 2). An important new result from this study, which is based on the first widespread sampling of the group, is that the South American species indeed form a distinct clade, the Neobartsia clade, that is very well supported with a posterior probability (PP) of 1.0 and a bootstrap support (BS) of 100. *Bellardia*–including the Neobartsia clade–is sister to *Odontites* (PP 1.0, BS 92) and together are sister to a clade comprised by *Hedbergia* and *Tozzia alpina* (PP 1.0, BS 93). The placement of the genus *Tozzia* was uncertain until now, although the support of our analyses is marginal (PP 0.94, BS 80). Finally, the genus *Euphrasia* is sister to the latter genera (PP 1, BS 100) and together form a clade sister to *Bartsia alpina* (PP 1, BS 100). This last clade is what Scheunert et al. (2012) referred to as the core Rhinantheae.

**Table 2.** Results for the Approximately Unbiased (AU) and the Shimodaira–Hasegawa (SH) tests at *p* < 0.05 for different constrained relationships. Log likelihood scores for the original analysis are given, as well as the difference in log likelihood between the original and the constraint topology (∂). Values in bold are significant with 95% confidence.

| nrDNA analysis constraint compared to clades from the<br>cpDNA analysis | ln likelihood | $\partial$ | AU | SH |
| --- | --- | --- | --- | --- |
| Unconstrained nrDNA analysis | −8445.56 |  |  |  |
| *Tozzia* + *Hedbergia* | −8449.04 | 3.48 | 0.166 | 0.186 |
| *H. decurva* + *H. abyssinica* var. *nykiensis* | −8464.51 | 18.95 | **0.004** | **0.026** |
| *Odontites* + *Bellardia* | −8456.43 | 10.87 | 0.141 | 0.131 |
| cpDNA analysis constraint compared to clades from<br>the nrDNA analysis |  |  |  |  |
| Unconstrained cpDNA analysis | −10044.47 |  |  |  |
| *Tozzia* + *Bellardia* | −10064.45 | 19.98 | **0.001** | **0.017** |
| *H. decurva* + *H. longiflora* ssp. *longiflora* | −10053.20 | 8.73 | 0.07 | 0.101 |
| *Odontites* + *Euphrasia* | −10071.61 | 27.14 | **0.0004** | **0.008** |

**Figure 1.**
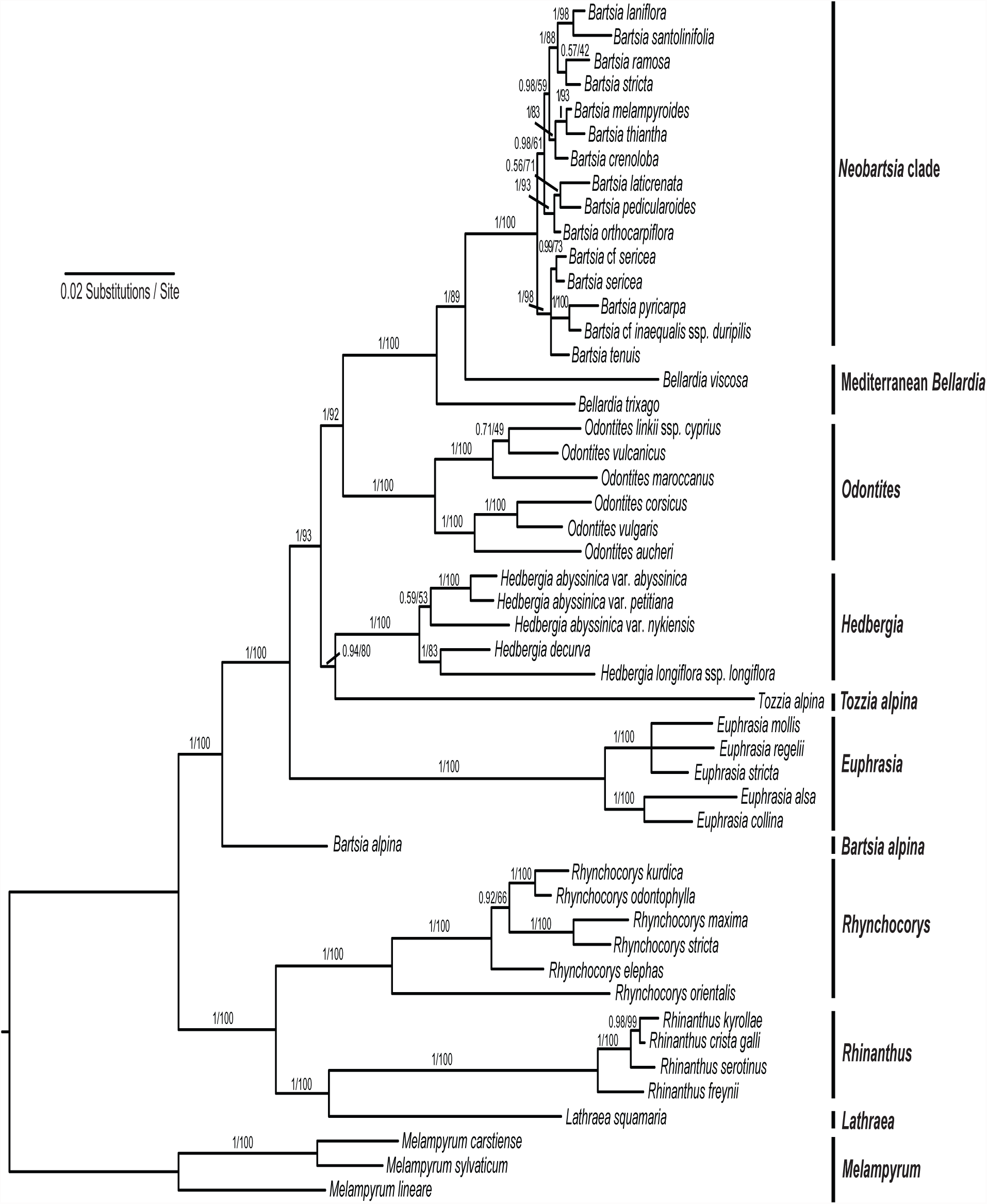
Majority rule consensus tree (excluding burn–in trees) with mean branch lengths from the partitioned Bayesian analysis of the combined dataset. Branch lengths are proportional to the number of substitutions per site as measured by the scale bar. Values above the branches represent Bayesian posterior probabilities (PP) and maximum likelihood bootstrap support (BS). Major clades are summarized following species names with the current species diversity in parenthesis.

**Figure 2.**
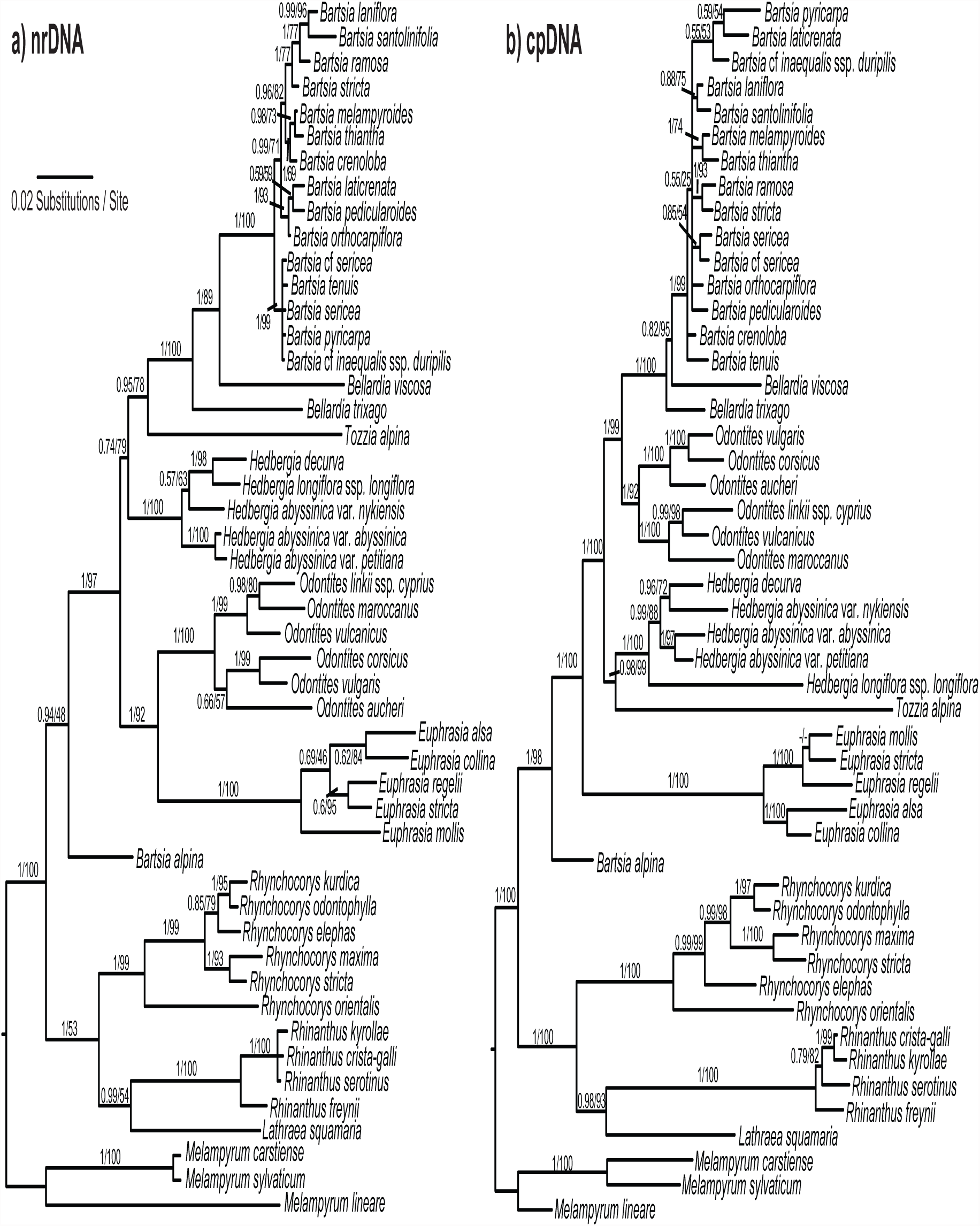
Majority rule consensus tree (excluding burn–in trees) with mean branch lengths from the partitioned Bayesian analysis of the a) nuclear ribosomal (nr) DNA and b) the chloroplast (cp) DNA datasets. Branch lengths are proportional to the number of substitutions per site as measured by the scale bar. Values above the branches represent Bayesian posterior probabilities (PP) and maximum likelihood bootstrap support (BS).

### Divergence Time Estimation

When the calibration point was placed at the node where *Melampyrum* diverged from the remaining genera, the South American Neobartsia clade had a median age of 2.59 Ma (1.51–4.08 Ma 95% HPD) (Table 3). The split between *Bellardia trixago* and the remaining species in the clade was estimated to have a median age of 8.73 Ma (5.12–12.76 Ma). The African clade diverged from *Tozzia alpina* 13.64 Ma (8.78–18.70 Ma), while the split of the European *Bartsia alpina* occurred 22.62 Ma (17.49–28.07 Ma). The root of the tree was estimated to have a median age of 30.65 Ma (25.55–38.83 Ma). Likewise, when the geological constraint was imposed, the Neobartsia clade had a median age of 2.63 Ma (1.97–3.58 Ma), and the divergence of *Bellardia trixago* from the remaining *Bellardia*–Neobartsia clade species occurred 8.48 Ma (4.95–12.48 Ma). The African clade diverged from *Tozzia alpina* 13.51 Ma (8.69–18.75 Ma), *Bartsia alpina* of 22.33 Ma (16.23–28.36 Ma), and the root of the tree was estimated at 30.98 Ma (29.13–35.96 Ma). The age of the root is consistent with the date (35.5 Ma) inferred for this clade in an angiosperm wide analysis (Zanne et al., 2014). Because the results using different calibration strategies were within the 95 percent HPD of each other (see Table 3), we used the root calibration analysis for subsequent biogeographic and diversification rate analyses.

**Table 3.**
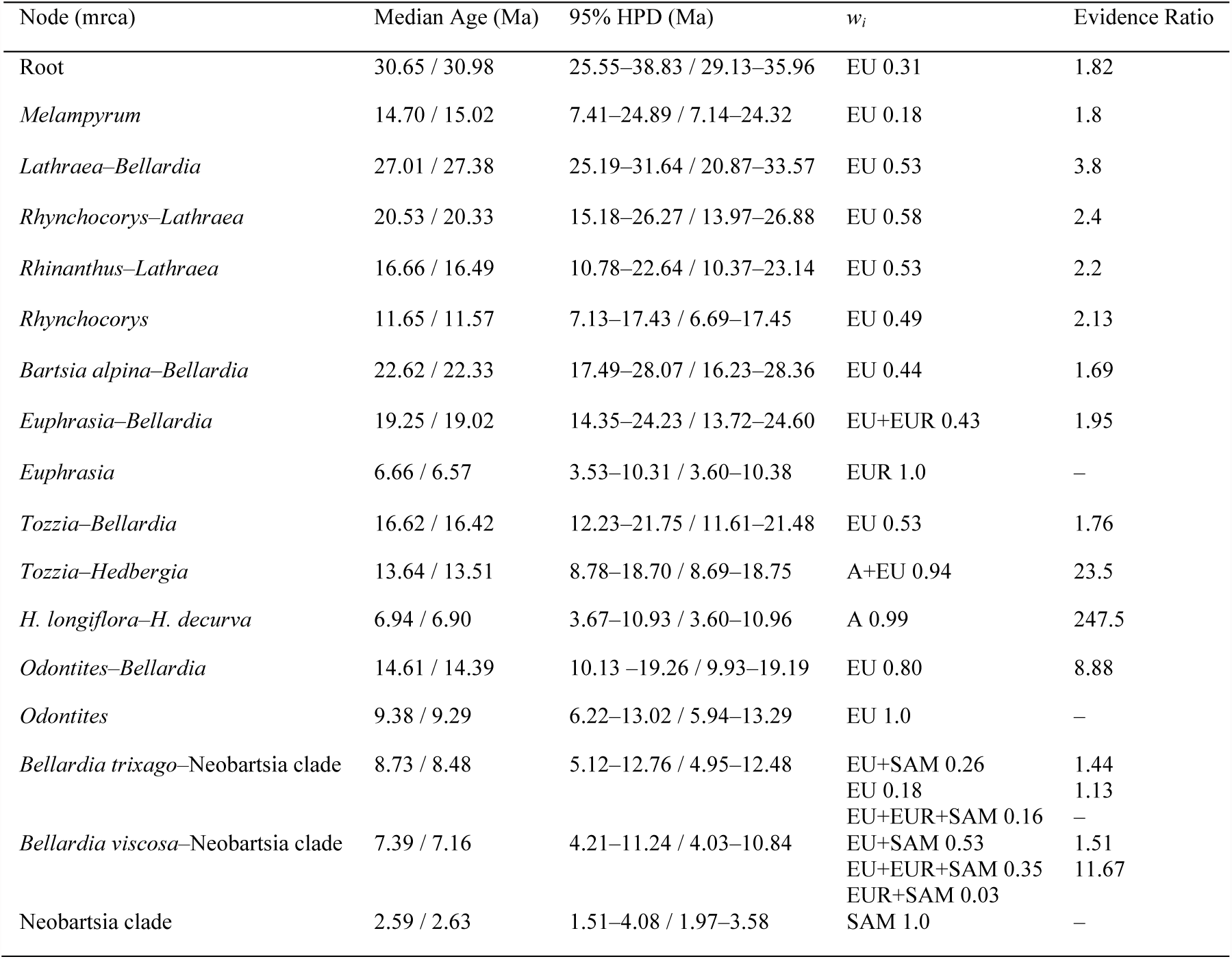
Divergence time estimates for the main clades, with each node representing the most recent common ancestor (mrca) of the taxa mentioned. The first value was obtained by calibrating the node of divergence of *Melampyrum* from its sister clade. The calibration point had a prior with lognormal distribution, offset 25 Myr, mean of 0.9, and standard deviation of 0.8, using the results of Wolfe et al. (2005), but incorporating considerable temporal uncertainty. The second value corresponds to an additional analysis where the uplift of the Andes was used as the calibration point of the node of divergence for S. Am. *Bellardia*. This last calibration had a prior with a lognormal distribution, offset of 1.7 Myr, mean of 0.2, and standard deviation of 0.6. Median age estimates as well as the 95% highest posterior density (HPD) are shown for both analyses. In addition, the composite Akaike weights (*w*_*i*_) from our biogeographic analyses are shown for the ‘expert based’ coding of current geographic distributions with the following abbreviations: A (Africa), EU (Europe), EUR (Eurasia), ENA (Eastern North America), SAM (South America). Evidence ratio is presented for the most supported geographic reconstruction.

### Biogeographic Analyses

Our three different codings of current geographic distribution resulted in very similar ancestral reconstructions (Table 4). Given that so much work has been done in recent years for several of these groups, e.g., *Bartsia/Bellardia* (Molau, 1990), *Euphrasia* (Gussarova et al., 2008), and *Odontites* (Bolliger, 1996), we favored the second coding scenario where current distributions were based on expert knowledge, including recent phylogenetic and biogeographic studies (for a wide-scale example on campanulids see Beaulieu et al., 2013). The most recent common ancestor (mrca) of the Rhinantheae clade of Orobanchaceae was likely distributed in Europe with a composite Akaike weight (*w*_*i*_) of 0.31 and an evidence ratio of 1.82 (Table 3). This ancestral range is maintained throughout the backbone of the tree until the node where *Euphrasia* diverges from the rest of the genera (*w*_*i*_ = 0.43, evidence ratio = 1.95). Nevertheless, a European ancestral range becomes the most supported reconstruction again at the node of divergence of *Odontites* (*w*_*i*_ = 0.80, evidence ratio = 8.88). A South American ancestral range is included for the first time at the crown node of *Bellardia*, where a *w*_*i*_ 0.26 supports a split between Europe and South America and a *w*_*i*_ of 0.18 supports an entirely European ancestral range; the evidence ratio between these two reconstructions is 1.44. A Third scenario supporting a range comprised of Europe, Eurasia, and South America is supported by a *w*_*i*_ of 0.16; the evidence ratio between this and the second most supported scenario is 1.13. The node where the Neobartsia clade diverges from the Mediterranean *Bellardia viscosa* is again supported by three competing models i) a split between Europe and South America (*w*_*i*_ = 0.53, evidence ratio = 1.51), ii) one between South America and an area comprised of Europe and Eurasia (*w*_*i*_ = 0.35, evidence ration = 11.67), and iii) one containing Eurasia and South America (*w*_*i*_ = 0.03). Additional results for other genera can be seen on figure 3 and summarized in table 4.

**Table 4.**
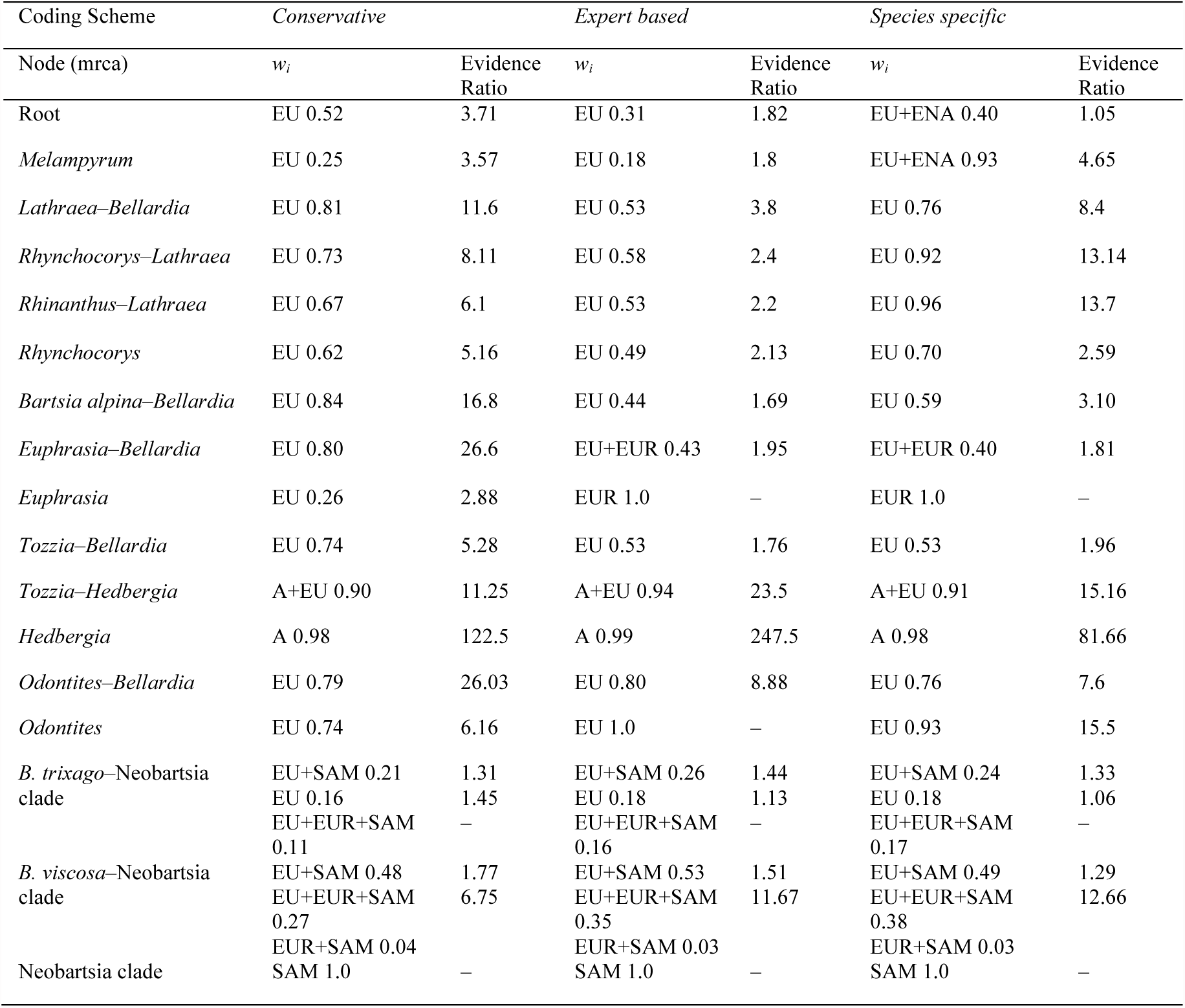
The composite Akaike weights (*w*_*i*_) are shown for our three different coding scenarios: conservative, expert based, and species specific. Abbreviations are as follow: A (Africa), EU (Europe), EUR (Eurasia), ENA (Eastern North America), SAM (South America). Evidence ratio is presented for the most supported geographic reconstruction.

**Figure 3.**
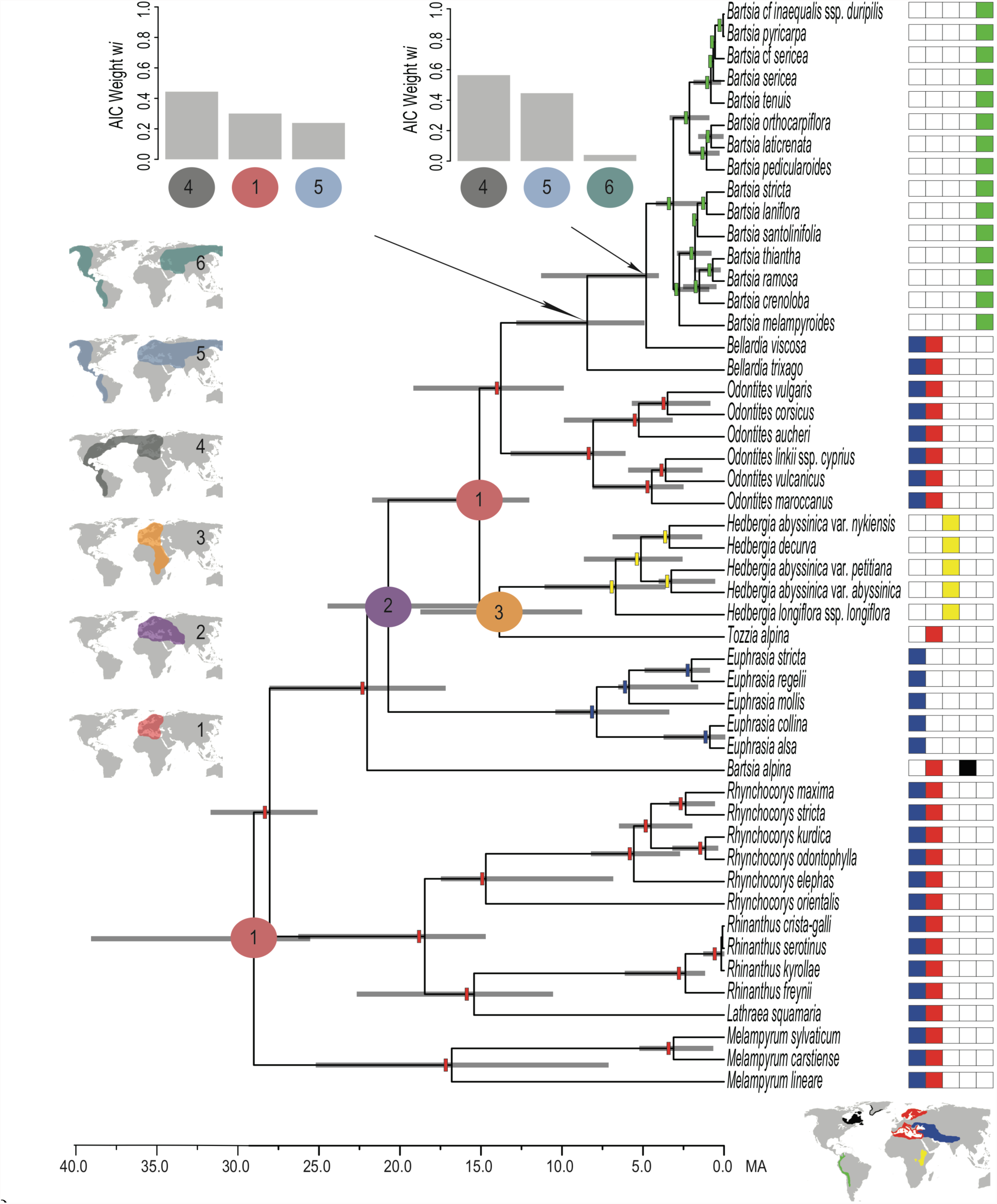
Topology obtained after combining and annotating five independent BEAST analyses. The calibration point was set at the node where all genera are included except for *Melampyrum*. The calibration had a prior with a lognormal distribution, offset 25 Ma, a mean of 0.9, and a standard deviation of 0.8 following dates by by Wolfe et al. (2005). Time in millions of years ago (Ma) is represented by the scale below the tree. Current distributions of the species are color–coded after the species names. The current distributions are plotted on a map below the species names and correspond to blue for Eurasia, red for Europe, yellow for Africa, black for northeastern North America, and green for South America. The most supported ancestral range reconstructions obtained from a Lagrange analysis, are plotted on the tree with color rectangles or circles with numbers that represent different biogeographic hypotheses. Informally drawn ancestral range reconstruction scenarios are plotted on five different maps on the left of the figure, each with a number that distinguishes it. Composite Akaike weights (*w*_*i*_) are plotted in the form of histograms for nodes where the reconstruction had competing hypotheses. Two possible routes of migration, one including the North Atlantic Land Bridge (NALB) and one including the Bering Strait, are shown on maps 5 and 6.

### Diversification Rates

The rates reported in this study correspond to different parts of the tree, known as breakpoints (e.g., nodes, stems, or both) that are evolving under different parameter values, i.e. per-lineage net diversification and relative extinction rates (Pennell et al., 2014). Our analyses discovered six shifts in the rate of net diversification (*r* = speciation minus extinction) in the Rhinantheae clade when performed over the posterior distribution of trees; three of these were also identified on the MCC tree. Importantly, the three shifts identified on the MCC tree corresponded to the shifts that occurred at the highest frequency in the analyses across the posterior distribution of trees. Because most of these analyses were conducted on a posterior distribution of trees to incorporate phylogenetic uncertainty (both temporal and topological), we report the mean net diversification rate of each shift (*r*_*mean*_) in the text, and the ranges of these shifts in table 5. For the three shifts found in the MCC tree, we also report that value (*r*_*mcc*_). The first two shifts found in our analyses correspond to shifts that were only present in less than 15 percent of the trees and show minimal deviation from the background rate of the tree. One of these shifts is on the node subtending the core Rhinantheae (*r*_*mean*_ = 0.11, frequency = 0.07) and the other one involves the hemiparasitic genus *Rhinanthus* L. and the holoparasite *Lathraea* L. (*r*_*mean*_ = 0.17, frequency = 0.12). The latter shift could correspond to a change in life history from hemiparasitism to holoparasitism in *Lathraea*, but given the limited sampling of these two groups and the low frequency at which the shift was found we dare not comment further. The next shift involves a slowdown in the rate of *Bartsia alpina* (*r*_*mean*_ = –0.4, *r*_*mcc*_ = 0) and was the most frequent shift in the analyses (frequency = 1.17). The frequency higher than 1.0 for this node is an artifact of the way MEDUSA adds the shifts. When two sister clades each have a shift at their crown nodes, MEDUSA adds the parameters from both shifts and places the result on the stem leading to the two clades. Thus these shifts do not occur with a frequency higher than 1.0, but are very common. The fourth shift corresponds to an increase in net diversification (*r*_*mean*_ = 0.09) in the clade sister to *Bartsia alpina* and was found with a frequency of 0.32. An additional shift was found in the clade comprised of *Tozzia alpina* and the genus *Hedbergia*, the shift was found in 75% of the trees (*r*_*mean*_ = –0.06; *r*_*mcc*_ = 0.05). Finally, a shift showing an uptick in net diversification rate was present for the Neobartsia clade, with a frequency of 0.40 (*r*_*mean*_ = 0.40; *r*_*mcc*_ = 0.79).

**Table 5.**
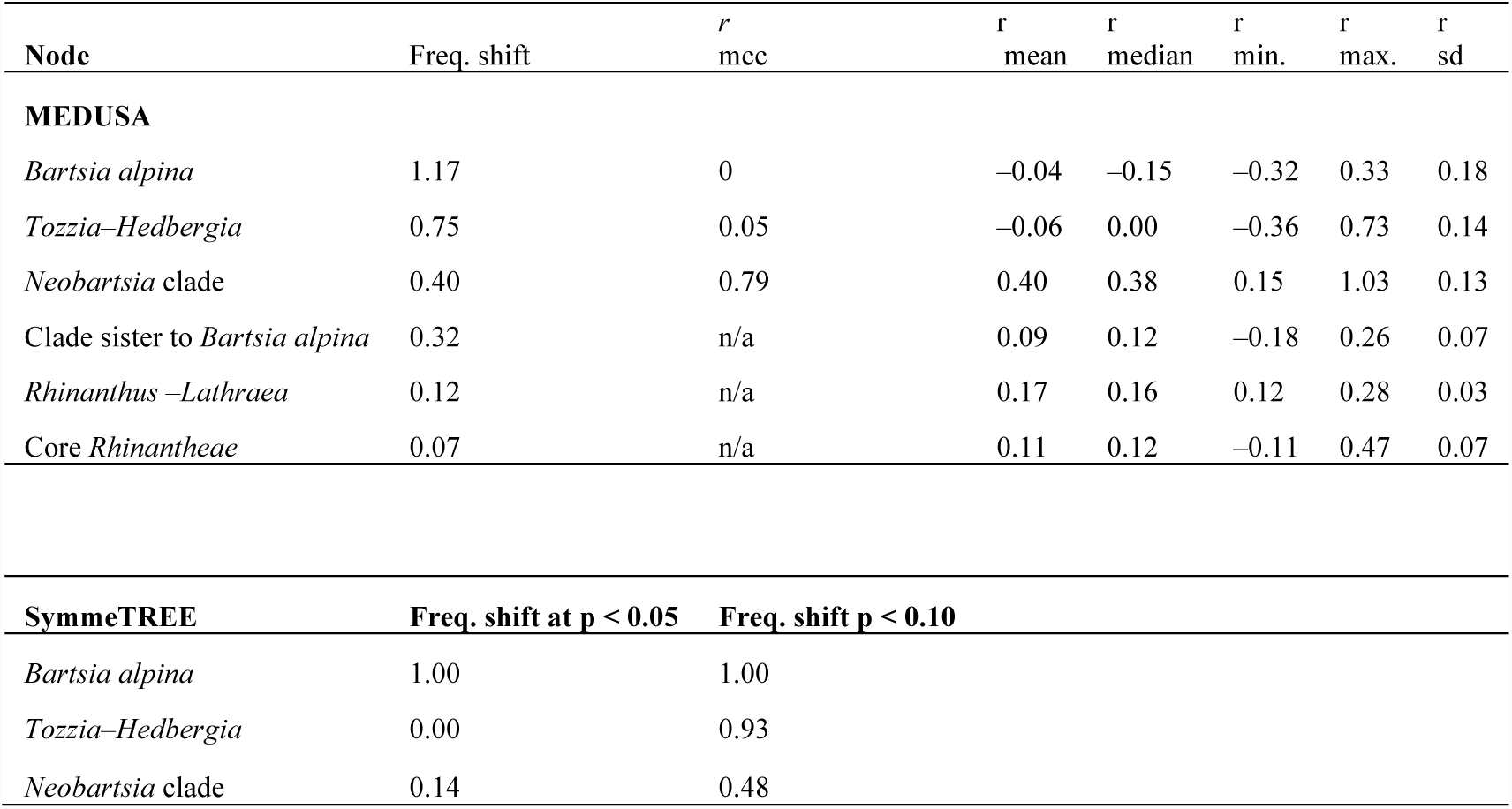
Results from our diversification rate analyses using MEDUSA and SymmeTREE over a posterior distribution of trees. The shifts were found at the nodes subtending the taxa specified in the first column, followed by the frequency of that shift in the posterior distribution of trees, the net diversification rate (*r*) for the maximum clade credibility tree (mcc), and the mean, median, minimum (min), maximum (max), and standard deviation (sd) summarized across 1,000 trees from the posterior distribution. In the results for SymmeTREE, two different significance values (α) were examined, α < 0.05, and α < 0.10.

| Node | Freq. shift | $r$<br>mcc | $r$<br>mean | $r$<br>median | $r$<br>min. | $r$<br>max. | $r$<br>sd |
| --- | --- | --- | --- | --- | --- | --- | --- |
| **MEDUSA** |  |  |  |  |  |  |  |
| *Bartsia alpina* | 1.17 | 0 | -0.04 | -0.15 | -0.32 | 0.33 | 0.18 |
| *Tozzia-Hedbergia* | 0.75 | 0.05 | -0.06 | 0.00 | -0.36 | 0.73 | 0.14 |
| Neobartsia clade | 0.40 | 0.79 | 0.40 | 0.38 | 0.15 | 1.03 | 0.13 |
| Clade sister to *Bartsia alpina* | 0.32 | n/a | 0.09 | 0.12 | -0.18 | 0.26 | 0.07 |
| *Rhinanthus-Lathraea* | 0.12 | n/a | 0.17 | 0.16 | 0.12 | 0.28 | 0.03 |
| Core *Rhinantheae* | 0.07 | n/a | 0.11 | 0.12 | -0.11 | 0.47 | 0.07 |
| **SymmeTREE** |  |  |  |  |  |  |  |
| | Freq. shift at $p < 0.05$ | Freq. shift $p < 0.10$ | | | | | |
| *Bartsia alpina* | 1.00 | 1.00 |  |  |  |  |  |
| *Tozzia-Hedbergia* | 0.00 | 0.93 |  |  |  |  |  |
| Neobartsia clade | 0.14 | 0.48 |  |  |  |  |  |

In comparison, the results obtained with SymmeTREE evidenced fewer diversification shifts on the whole tree (*p* < 0.05). Like the MEDUSA results, an increase in diversification rate was also leading to the clade sister to *Bartsia alpina* and was consistently found in every tree we analyzed (Table 5). A shift showing a slowdown in the *Tozzia+Hedbergia* clade was found to be marginally significant at p < 0.05 (p = 0.067) in every tree. If we were to choose a less stringent significance threshold (e.g., p < 0.10), this shift would be significant in 506 trees (93% of the distribution). The same is true for the shift involving the Neobartsia clade, where it was only found to be significant in 77 trees at *p* < 0.05 (14%), but increased to 258 trees (48%) when the less stringent *p* value was chosen.

## DISCUSSION

### Systematic Implications

Molau (1990) published a comprehensive monograph on the genus “*Bartsia*”, where he hypothesized that the species formed a monophyletic group that was sister to the African monotypic genus *Hedbergia.* Our phylogenetic results (Figs. 1 and 2), which are in agreement with those of Těšitel et al. (2010) and Scheunert et al. (2012), show clearly that *Bartsia* sensu Molau is polyphyletic, and that the new classification (sensu Scheunert et al., 2012) better reflects the disparate geographic distributions of these lineages, as well as their evolutionary histories. Previous studies have recovered the basal relationships within *Bellardia* as a polytomy (Těšitel et al., 2010; Scheunert et al., 2012), where the position of *B. trixago, B. viscosa*, and *B. latifolia* is uncertain. Here, we recovered *Bellardia trixago* as sister to *B. viscosa* and the Neobartsia clade, but because we did not sample *B. latifolia*, we cannot be certain of the position of the other two taxa. The South American Neobartsia clade was highly supported in every analysis, and this is the first study to sample a geographically and morphologically representative diversity of the richness in this clade. These results provide strong evidence of the evolutionary distinctiveness of the Neobartsia clade with respect to the Mediterranean members of the expanded genus *Bellardia* (sensu Scheunert et al. 2012) – i.e., its unique geographic distribution and biogeographic history, the long divergence times from their Mediterranean relatives (∼7.39 Ma), and the elevated diversification rates. Along with diagnostic morphological characters, we feel this justifies a reanalysis of the generic revision of Scheunert et al. (2012) with respect to the taxonomy in this clade, and this is the subject of ongoing taxonomic work in this clade.

Our cpDNA and nrDNA analyses placed the genus *Tozzia* in different positions in the tree, although these differences were not statistically significant (Table 2). Our combined analysis placed the genus as sister to *Hedbergia*, albeit with marginal support (PP 0.94, BS 80). While this relationship is in agreement with previous studies (Těšitel et al., 2010; Scheunert et al., 2012), further work will be necessary to confidently place this genus in the Rhinantheae clade. Lastly, the African genus *Hedbergia* showed interesting and likely problematic species delimitations. The taxon *H. abyssinica* var. *nykiensis* was sister to *H. decurva* in our cpDNA analyses, and its relationship to the other *H. abyssinica* varieties in the nrDNA dataset was weakly supported. This is the first time that varietal taxa for this group have been included in a molecular study, and highlights the necessity for a more detailed study on the clade.

### Biogeography and Diversification Rates

Our divergence time and biogeographic results depicted in figure 3 illustrate evolutionary hypotheses regarding the current distribution of the Rhinantheae clade. As a reminder, our analyses were calibrated using dates obtained with the molecular rate of the ITS (Wolfe et al., 2005), and thus, they should be taken as estimates where some uncertainty is expected. Nevertheless, they provide an evolutionary foundation that helps explain the current distribution and the diversity of the South American Neobartsia clade. With no doubt, Europe played a major role, almost at every node, in the reconstruction of ancestral ranges in the Rhinantheae clade of Orobanchaceae. Although this is the first formal biogeographic analysis in the clade, these results are in line with the verbal biogeographic scenarios described in previous studies (Wolfe et al., 2005; Těšitel et al., 2010), but with slight differences in the description of the ancestral areas. Diversification of the majority of the genera was achieved in the European continent with subsequent migration events to Eurasia, northeastern North America, the Mediterranean region, Africa and South America. *Bartsia alpina* is a good example of a taxon with a purely European ancestral distribution but that is currently distributed in other parts of the world. This suggests that the current distribution was the result of a second and more recent migration into Greenland and northeastern North America sometime along its very long branch (Fig. 3). *Odontites* is another good example of a European radiation that has expanded its range to include Eurasia after the initial divergence. Moreover, the genus *Euphrasia*, which accounts for more than half of the members of the clade with ∼400 species, was reconstructed as having a Europe/Eurasian ancestral range. The few species sampled in this study all have Eurasian distributions but the genus is currently considered to have a “bipolar” distribution (Gussarova et al., 2008), with species distributed in north temperate regions and extreme Austral areas. This pattern is extremely interesting since it suggests that extinction and/or long distance dispersal have played a large role in shaping the current distribution of this large clade.

The Mediterranean region was not included as a distinct area in our reconstruction analyses, and therefore, some of the genera with current Mediterranean distributions were treated as European, e.g., *Rhynchocorys, Odontites*, *Bellardia trixago*, and *B. viscosa*. The Mediterranean climate, as recognized today, is a young environment formed only 2.3–3.2 Ma and it is the result of two main events: i) the establishment of the Mediterranean rhythm of dry summers and mild–cold winters ∼3.2 Ma, and ii) the oldest xeric period know for the region ∼2.2 Ma (Zagwin, 1960; 1974; Suc, 1984). The crown clades for each of these genera were reconstructed to have a European ancestral distribution, which implies that their current ranges are the results of independent evolutions into the European Mediterranean climatic region not earlier than ∼3.2 Ma.

This study is mainly focused on studying the diversity of the Neobartsia clade in the Andes, and to propose plausible hypotheses for its distribution. The Andes are thought to have begun uplifting in the late Miocene (∼10 Ma) but only reaching the necessary elevation to host alpine conditions in the late Pliocene or early Pleistocene 2–4 Ma (Simpson, 1975; Burnham and Graham, 1999; Gregory-Wodzicki, 2000; Antonelli et al., 2009). In our biogeographic analyses, South America is reconstructed for the first time at the crown node of *Bellardia*, with a median age of 8.73 Ma (5.12-12.76 Ma), and then at the node where *Bellardia viscosa* and the Neobartsia clade diverge (median age of 7.39 Ma [4.21-11.24 Ma]).

These reconstructions, between 12.76 and 4.21 Ma, define an eight and a half million year window for the ancestor to have reached South America. There are two main land routes that were present during this time period, the North Atlantic Land Bridge (NALB) uniting northeastern North America and western Europe, and the Bering Land Bridge between eastern Asia and western North America. Previous studies in the plant family Malpighiaceae (Davis et al., 2002; 2004), have suggested a migration route from South America to Africa starting in the early Oligocene (∼30 Ma) via North America, the NALB, and Europe. The NALB was available from the early Eocene (∼50 Ma) until the middle to late Miocene (∼10–8 Ma) (Tiffney, 1985; Tiffney and Manchester, 2001; Denk et al., 2010; 2011), dates which overlap with our divergence time estimates (Table 3) and with the appearance of South America as an ancestral range in our biogeographic analyses. This allows for the possibility of an early dispersal from Europe into North America over this land bridge. Colonization of North America would have followed a stepwise migration to South America over the forming Isthmus of Panama and/or island chains sometime in the last 4.5 Ma (Coates et al., 2004; Kirby and MacFadden, 2005; Retallack and Kirby, 2007). A caveat of this scenario is that it leaves a narrower window of time for a stepwise migration to have occurred. Moreover, the warmer temperatures in eastern North America during the late Miocene would possibly have affected the migration of a presumably alpine adapted ancestor through the NALB.

An alternative stepwise migration scenario for the South American clade’s colonization of the Andes involves a migration route through Beringia. This land bridge, which was available on–and–off from ∼58–3.5 Ma (Hopkins, 1967; Tiffney and Manchester, 2001; Tiffney, 2008), has been proposed as a route for several groups found both in eastern Asia, western north America, and the Andes—e.g., Valerianaceae (Moore and Donoghue, 2007). The age of this land bridge overlaps completely with both the divergence of the Neobartsia clade from *Bellardia viscosa* (4.21-11.24 Ma), as well as with the split between *Bellardia trixago* and the other members of the *Bellardia*–Neobartsia clade 5.12–12.76 Ma. Moreover, this more recent route allows for the world to cool down during the Pliocene (Tiffney and Manchester, 2001), which may have facilitated the migration. This migration scenario is also plausible since Eurasia, Europe, and South America were reconstructed as the second most supported ancestral range at the node of divergence of the South American clade (*w*_*i*_ = 0.35). Both of these stepwise migration scenarios rely completely on North America as an intermediary step where the South American ancestor possibly diversified, migrated, and finally went extinct. Unfortunately, there is no fossil record in the Rhinantheae clade (or in Orobanchaceae), and thus, no physical evidence is available to support either of these hypotheses.

Molau (1990) hypothesized that the Neobartsia clade had colonized the Andes via a long–distance dispersal from Africa, sometime in the early Pliocene (∼5 Ma). This hypothesis seemed plausible at the time when no phylogenetic evidence was available for the clade, but now that it is clear that the former genus *Bartsia* is polyphyletic and the two African species (*Hebergia decurva, H. longifolia*) are not sister to the South American species, there is no longer support for this hypothesis. Nevertheless, there is a third hypothesis that does rely on long–distance dispersal, but rather from Mediterranean Europe/north Africa to Andean South America (or, alternatively, from somewhere in North America following a land bridge migration from the Old World). Many plants are dispersed over long distances by water (e.g., *Cocos* L.), birds (e.g., *Pisonia* L.), or wind (e.g., *Taraxacum* F.H. Wigg.), and physiological and morphological adaptations to float, adhere, or fly are common (reviewed in Howe and Smallwood, 1982). The seeds of *Bellarida*–Neobartsia clade are enclosed in a dry dehiscent capsule that contains between 20–200 small seeds (0.3–2 mm) per fruit, each equipped with 6–13 short wings or ridges (Molau, 1990). Although these seeds are light and have wings making them at first glance suitable for long distance traveling, it has been estimated that their mean dispersal distance is 0.3 meters, at least in *Bartsia alpina* (Molau, 1990). The short mean dispersal distance is in strong disagreement with the distance that a seed would need to travel from the Mediterranean region to the New World (∼7,000 km/∼4,000 mi). Nevertheless, there is a known constant storm track from western Africa (including the northwestern African Mediterranean climatic region) that crosses the Atlantic Ocean into the Caribbean and the Americas, and recent evidence has shown that there are major influxes of African dust in southern North America (Bozlaker et al., 2013), northeastern South America (Prospero et al., 2014), and the Caribbean basin (Prospero and Mayol-Bracero, 2013). This opens the possibility for seeds of a Mediterranean ancestor to have been picked up and carried over to the New World. Although at first this may seem unlikely, it is important to point out that a single seed may be sufficient for the colonization of a new habitat, and that the eight and a half million year time window coupled with the large amounts (∼200) of seeds that are produced in each capsule, increase the probability for this event to have happened.

At this point we cannot accept or reject any of the biogeographic hypotheses described above—the two stepwise migrations through North America or the long–distance dispersal from the Mediterranean climatic region—and it highlights the difficulty of inferring ancestral colonization routes even when using modern ancestral range reconstruction methods (see Tripp and McDade, 2014), especially with a non–existent fossil record.

To investigate if these biogeographic movements have affected the rate at which clades are diversifying (i.e., “dispersification”), we need to assess if the shifts found in our analyses correlate with a movement into a new area or if there is something else, e.g., a morphological change, that has triggered them. Regardless of the reason, investigating shifts of diversification and the location of these on a phylogenetic tree is extremely helpful when trying to understand disparities in species richnesses across related clades. The comparison of two methods that are based on different tenets, a stepwise model testing approach vs. a topological imbalance approach (MEDUSA and SymmeTREE, respectively), allowed us to i) better evaluate the performance of different approaches used to identify shifts in diversification, while ii) making results shared by both methods robust and reliable. This comparison also showed the advantages of using a stepwise model testing approach and a method that incorporates extinction. Our MEDUSA analysis found six shifts across the posterior, and three when using the MCC tree; two of these six identified shifts represent a slowdown in net diversification. One of these slowdowns, which is the only shift consistently found by SymmeTREE at *p* < 0.01, across the posterior, and in the MCC tree, corresponds to the node where *Bartsia alpina* diverges from the rest of the core Rhinantheae 22.62 Ma. The extremely low diversification rate and its very long branch indicate that this species is likely the only extant member of a lineage that has had historically very low speciation rates or high extinction rates, or both. The first significant increase in net diversification rates was found at the node where the genus *Euphrasia* diverges from other genera 19.25 Ma. This genus includes ∼400 species that encompass more than 80% of the species richness of Rhinantheae, estimated to be ∼528 spp. (Mabberley, 2008). Based on our limited sampling of this group, we cannot identify an apparent change in morphology or geography in the genus, and thus, no evident cause for this shift can be assessed with these data. Nevertheless, given the age and very high diversity of the clade, this shift is not surprising. However, it is important to point out that because we collapsed clades at the generic level to incorporate unsampled diversities, the present shift might not be the only one in *Euphrasia* and that clades within the genus may also have shifts of their own, where there might be an apparent change in either morphology or geography.

We also identified an increased rate of net diversification in the South American Neobartsia clade. We hypothesize that the clade underwent a similar pattern as seen in other Andean radiations, e.g. the family Valerianaceae and the genus *Lupinus* (Bell and Donoghue, 2005b; Hughes and Eastwood, 2006, respectively), where their North American ancestor was “pre–adapted” to cold environments making the colonization of the high Andes, and further radiation, easier (Donoghue 2008). The Neobartsia clade has a median divergence time of 2.59 Ma and a mean diversification rate of 0.40, however, when the analysis is performed on the MCC tree, the net diversification almost doubles (*r*_*mcc*_ = 0.79). The large difference in values implies that although the shift was only identified in 40% of the posterior distribution of trees, when detected, the rate can be nearly four times higher than the background rate of the tree (background *r*_*mean*_ = 0.22). Based on the very short branches within the clade, its young age, and the genetic similarity between the species included in this study, this shift likely resulted in a rapid radiation event where the movement to and colonization of the high Andes acted as a trigger to increased diversification. As the Andes were uplifting, the creation of alpine conditions promoted the radiation into the diversity that we see today. Accordingly, this is a another example of how the movement into a new geographic area, can lead to a high number of species in a relatively short period of time without the appearance of morphological key innovation, which is what Moore and Donoghue (2007) referred to as “dispersification”.

### Conclusions

This study places the Neobartsia clade in the context of a robust and well–supported phylogeny within the Rhinantheae clade of Orobanchaceae. This is the first study to study this clade in an explicitly temporal framework, with detailed divergence time estimates for the clade. Here, we focused primarily on the colonization and diversification of Andean South America ∼2.59 Ma. This date correlates well with the necessary age for the Andes to have acquired the adequate elevation to simulate alpine conditions for the establishment of this temperate, largely alpine clade in South America. Given that the South American clade is sister to a Mediterranean taxon, we hypothesized three biogeographic scenarios for the colonization of the Andes. The first route involves the NALB and North America as a stepwise migration route from Europe ∼12–8 Ma, whereas the second hypothesis involves a westerly route from Europe through Asia, the Bering Land Bridge, and North America ∼12–4 Ma. Both of these scenarios share a second migration from North America to South America over the forming Isthmus of Panama and/or island chains in the mid to late Pliocene ∼4.5–3.13 Ma, which gave rise to the Neobartsia clade, and high levels of extinction throughout Asia and/or North American. Finally, the third hypothesis involves a long–distance dispersal from the Mediterranean climatic region (Europe and northern Africa) to South America. At this point however, we cannot accept or reject any of the previously described hypotheses. Regardless of the biogeographic route taken, once the South American ancestor reached the Andes, it was able to diversify rapidly in the vacant niches in the páramos. The greater diversification rates in the Neobartsia clade help explain the species richness found in the Andes today and support the idea that the “key opportunity” of geographic movement into a new area may trigger high diversification without the necessity of the evolution of morphological key innovations, and this may be especially true when the colonizing ancestral lineage is adapted to the new conditions it encounters.

